# A Rapid, Functional sgRNA Screening Method for Generating Murine RET and NTRK1 Fusion Oncogenes

**DOI:** 10.1101/2023.04.06.535912

**Authors:** Laura Schubert, Anh T. Le, Trista K. Hinz, Andre Navarro, Sarah K. Nelson-Taylor, Raphael A. Nemenoff, Lynn E. Heasley, Robert C. Doebele

## Abstract

CRISPR/Cas9 gene editing technology is an indispensable and powerful tool in the field of cancer biology. To conduct successful CRISPR-based experiments, it is crucial that sgRNAs generate their designed alterations. Here, we describe a simple and efficient sgRNA screening method for validating sgRNAs that generate oncogenic gene rearrangements. We used IL3-independence in Ba/F3 cells as an assay to identify sgRNA pairs that generate fusion oncogenes involving the *Ret* and *Ntrk1* tyrosine kinases. We confirmed these rearrangements with PCR or RT-PCR as well as sequencing. Ba/F3 cells harboring *Ret* or *Ntrk1* rearrangements acquired sensitivity to RET and TRK inhibitors, respectively. Adenoviruses encoding Cas9 and sgRNAs that catalyze the *Kif5b-Ret* and *Trim24-Ret* rearrangements were intratracheally instilled into mice and yielded lung adenocarcinomas. A cell line (TR.1) was established from a *Trim24-Ret* positive tumor that exhibited high *in vitro* sensitivity to RET-specific TKIs. Moreover, orthotopic transplantation of TR.1 cells into the left lung yielded well-defined tumors that shrank in response to LOXO-292 treatment. The method offers an efficient means to validate sgRNAs that successfully target their intended loci for the generation of novel murine oncogene-driven tumor models.

## Introduction

Cancer genomes are characterized by numerous and often complex genetic alterations [1, 2]. Dissecting the functional role of each of these alterations can be challenging, but has been crucial to understanding fundamental underpinnings of cancer biology. Small molecule inhibitors are available for treating *ALK*, *ROS1, RET* and *NTRK1-*rearranged receptor tyrosine kinases (RTKs) in lung cancers, but variable or incomplete responses are often observed and acquired resistance is unavoidable [3-7]. To this end, human tumor-derived cell lines and patient-derived xenografts (PDXs) have been mainstays for the mechanistic and functional exploration of these genetic alterations as well as drug resistance mechanisms [8-10]. The pace of discovery of the oncogenic drivers in histologically-defined cancers such as lung adenocarcinoma is not always matched with access to human cell lines or PDXs as preclinical models. Moreover, a growing literature reveals that the tumor microenvironment (TME) including host immune cells significantly contribute to the therapeutic responses achieved with oncogene-targeted agents like tyrosine kinase inhibitors (TKIs) and KRAS^G12C^-targeted agents [11-17]. Thus, rigorous preclinical modeling of oncogene-targeted agents relevant to that observed in patients requires input from the host adaptive immune system. In this regard, experiments with existing human cell lines and PDXs performed in immune-deficient mice present significant limitations that can be overcome with murine models of oncogene-driven cancers permitting direct evaluation of cancer cell-TME interactions in fully immune-competent hosts.

The development of CRISPR/Cas9-based technologies allows precise manipulation of genes in their endogenous loci and has become an essential tool for modeling mutations and chromosomal abnormalities [18]. In this system, the Cas9 DNA endonuclease associates with a single guide RNA (sgRNA), thereby directing the complex to a specific locus of the genome via base complementarity. At this locus, Cas9 creates a double stranded break (DSB) upstream of a protospacer adjacent motif (PAM) site [19-23]. While CRISPR/Cas9 has been incredibly useful, it is well documented that there can be off-target effects or inefficient cleavage [24]. This limitation necessitates rigorous testing of sgRNAs to confirm that they generate their intended alterations. Often, multiple sgRNAs must be tested, which can be both time and labor intensive. Herein, we developed a CRISPR/Cas9 strategy to generate distinct RTK fusion oncogenes that are found across diverse cancer types including lung cancer [2]. These murine CRISPR-generated fusion oncogene models are predicted to greatly expand our ability to study mechanisms mediating therapeutic input from the TME and rapidly acquired drug resistance in syngeneic, immune-competent mouse models.

## Materials and Methods

### Ba/F3 cell culture and transfection

As previously described [25], Ba/F3 cells were cultured in RPMI1640 (Corning) supplemented with 10% fetal bovine serum (FBS) and 1 ng/mL IL3 (R&D Systems) where indicated. Cells were maintained in a humidified incubator at 37°C and 5% CO_2_.

The pMSCV-loxp-dsRED-loxp-eGFP-Puro-WPRE construct (a gift from Hans Clevers (Addgene plasmid #32702)) was stably introduced into Ba/F3 cells via retroviral transduction followed by puromycin selection. Fluorescence-activated cell (FAC)-sorting of single cell suspensions was used to select for dsRED+ cells after transduction. Cells were suspended in PBS with 1mM EDTA, 25mM HEPES (pH 7.0) and 1% FBS and sorted with the Astrios EQ (Beckman Coulter) into RPMI1640 (Corning) with 10% FBS at the CU Cancer Center Flow Cytometry Shared Resource. Ba/F3 cells were then transfected with 5 μg of total sgRNA plasmid (described below) via electroporation at 300V and 950μF using a BioRad GenePulser Xcell. Three days post-transfection, cells were suspended in PBS and analyzed for GFP and dsRED expression by flow cytometry on the YETI (Propel Labs) at the CU Cancer Center Flow Cytometry Shared Resource and analyzed using Kaluza Software (Beckman Coulter).

### Instillation of recombinant adenoviruses into murine lungs and establishment of the TR.1 murine ***Trim24-Ret* lung cancer cell line**

A recombinant adenovirus construct pAV-U6-Trim24-U6-Ret,CMV-hCas9:P2A:Cre (VectorBuilder, Chicago, IL) encoding Cas9, Cre recombinase and the *Trim24-Ret* sgRNA pair shown in **Suppl. Fig. 1** was packaged by ViraQuest, Inc (North Liberty, IA). C57BL/6 mice bearing floxed TP53 alleles (B6.129P2-Trp53^tm1Brn^/J; Jackson Laboratory stock #008462) were submitted to intratracheal instillation of 50 μL Adeno-TR-Cas9-Cre virus at a dose of 1.5 × 10^8^ PFU/mouse. Lungs from tumor-bearing mice 6 weeks after adenovirus-infection were harvested and tumors minced, digested and cultured in RPMI-1640 medium containing 5% fetal bovine serum until a stable epithelial cell line (TR.1) was established. The presence of the predicted *Trim24-Ret* fusion protein was confirmed by anti-Ret immunoblot analysis.

### Inhibitors and reagents

Ponatinib, cabozantinib and foretinib were purchased from SelleckChem (Houston, TX). Alectinib was obtained from Chugai Pharmaceuticals and entrectenib was obtained from Igynta. LOXO-292 and BLU-667 were purchased from MedChemExpress (Monmouth Junction, NJ). The RET antibody (EPR2871) was purchased from Abcam, pRET Y1062 (sc-20252) was purchased from Sana Cruz Biotechnology (Dallas, Texas) and GAPDH (65C) were purchased from Millipore.

### sgRNA design and cloning

The pX330-U6-Chimeric_BB-CBh-hSpCas9 plasmid (a gift from Feng Zhang (Addgene plasmid #42230)) was modified to express a bicistronic peptide containing Cas9 and Cre as follows. pX330 was sequentially modified to accept a 2.4kb Cas9-P2A-Cre fragment from pSECC (a gift from Tyler Jacks (Addgene plasmid # 60820) first by an EcoRI (New England Biolabs) digestion with a subsequent fill-in reaction by DNA polymerase and then a AccIII restriction digest. The 2.4kb insertion fragment was generated from pSECC by a restriction digest with AccIII (Promega) and SacII (New England Biolabs). Ligation of this fragment into the pX330 modified plasmid was performed using T4 DNA Ligase (Invitrogen) in an overnight reaction at 16°C. The ligated product was then used to transform competent bacteria. sgRNAs targeting *Trim24, Kif5b, Tpm3, Ret* and *Ntrk1* were designed using the Zhang lab CRISPR design tool (crispr.mit.edu) and are presented in **Supplementary Figure 1**. The pX330+Cre plasmid was digested with BbsI (New England BioLabs) and ligated to annealed and phosphorylated sgRNA oligonucleotides (Integrated DNA Technologies). Ligated plasmids were transformed into DH5α *E.coli* (Life Technologies). The PX330-*Alk-Eml4* plasmid was kindly provided by Andrea Ventura [26].

### Proliferation assays

Ba/F3 cells were plated (10,000 cells per well) in 96-well tissue culture plates in RPMI1640 (Invitrogen) supplemented with 10% FBS, with or without 1 ng/mL IL3 where indicated and treated with the indicated concentrations of drugs for 72 hours. CellTiter 96 MTS was used to estimate cell numbers according to the manufacturer’s instructions (Promega). Each assay was performed in triplicate with three biological replicates. For analysis of the sensitivity of the murine *Trim24-Ret* lung cancer cell line, TR.1, cells were seeded (200 per well) in 96-well plates and 24 hrs later, incubated for 7 days in triplicate with increasing doses (0-300 nM) of LOXO-292 or BLU-665. Cell number was assessed by measuring DNA content using the CyQUANT Direct Cell Proliferation Assay (Life Technologies, #C35011, Carlsbad, CA) according to manufacturer’s instructions. IC_50_ values were calculated using GraphPad Prism software.

### Immunoblotting

Immunoblotting was performed as previously described [8]. Briefly, cells were lysed in radioimmunoprecipitation assay buffer (RIPA) supplemented with Halt protease and phosphatase inhibitor (Thermo Scientific). Lysates were diluted in 4X protein sample loading buffer (LI-COR) and subjected to SDS-PAGE. Proteins were transferred to nitrocellulose and stained with primary antibodies followed by IRDye anti-mouse or anti-rabbit IgG (LI-COR). The Odyssey Imager and Odyssey Image Studio Software (LI-COR) were used to scan membranes and analyze images.

### DNA isolation, PCR and sequencing

DNA was isolated using the Quick-gDNA MiniPrep Kit from Zymo Research according to the manufacturer’s instructions. PCR was performed with primers designed to detect *Eml4-Alk, Eml4, Trim24-Ret, Tpm3-Ntrk1* and *Ret.* PCR products were separated with electrophoresis on a 1% agarose gel. Standard Sanger sequencing was performed on PCR-amplified fragments.

### RNA isolation, RT-PCR and sequencing

RNA was isolated using the RNEasy plus MiniKit (Qiagen) according to the manufacturer’s instructions. cDNA was generated using the SuperScript III First-Strand Synthesis System (Invitrogen) using random hexamers as per the manufacturer’s instructions. PCR reactions were performed with primers designed to detect *Kifb5-Ret* fusion transcripts. PCR products were separated with electrophoresis on a 1% agarose gel. PCR amplified-cDNA was subjected to standard Sanger sequencing.

### Orthotopic mouse model of RET lung cancer

The TR.1 cell line was propagated as orthotopic tumors in the left lung lobe of nine-week-old C57BL/6 mice (WT; C57BL/6J; #000664, Jackson Laboratory, Bar Harbor ME). Cells were prepared in a solution of 1.35 mg/mL Matrigel (Corning #354234) diluted in Dulbecco’s PBS (Corning) for injection. Mice were anesthetized with isoflurane, the left side of the mouse was shaved, and a 1 mm incision was made to visualize the ribs and left lobe of the lung. Using a 30-gauge needle, 2.5 × 10^5^ cells were injected in 40 µL of Matrigel-cell mixture directly into the left lobe of the lung and the incision was closed with staples. Tumors were permitted to establish for 10 days and then the mice were submitted to micro-computed tomography (µCT) imaging to obtain pre-treatment tumor volumes. Tumor-bearing mice were randomized into treatment groups (*n* = 5), either 10 mg/kg selpercatinib/LOXO-292 or diluent control (0.5% HPMC) by oral gavage 5 days/week until the end of study. Mice were imaged weekly by µCT imaging to monitor effects of drug treatment on tumor volume. Tumor volume µCT imaging was performed by the Small-Animal IGRT Core at the University of Colorado Anschutz Medical Campus in Aurora, CO using the Precision X-Ray X-Rad 225Cx Micro IGRT and SmART Systems (Precision X-Ray, Madison, CT). Tumor volume was quantified from µCT images using ITK-SNAP software<u>^36^</u> (www.itksnap.org). Upon study end mice were sacrificed using CO_2_ and cervical dislocation as a secondary method.

## Results

Using CRISPR to engineer chromosomal rearrangements encoding precise fusion oncogenes presents a unique challenge; two independent sgRNAs are necessary to generate double stranded breaks at both the 5’ and 3’ fusion partners. Previous studies have shown that CRISPR systems can be successfully used to generate both inter-and intra-chromosomal rearrangements *in vitro* [27-29]. Previously, this technology was used *in vivo* with either GEMMs or viral delivery methods [26, 30].

Typically, multiple pairs of sgRNAs must be tested to identify those that generate successful fusions. We sought to devise a strategy to quickly validate and enrich for sgRNAs that generate the intended rearrangement. The Ba/F3 murine B-cell line is dependent on exogenous IL3 for growth and proliferation, but can be rendered IL3-independent if they undergo an oncogenic transformation. We hypothesized that IL3-independence in Ba/F3 cells could be used to screen and select for sgRNAs that successfully generated oncogenic alterations. Briefly, Ba/F3 cells are co-transfected with a pair of sgRNA-encoding plasmids targeting the 5’ and 3’ fusion partners, IL3 is removed and cells are cultured until IL3-independent clones emerge (**Figure 1A**). Natural selective pressures enrich for productive sgRNAs pairs, thereby eliminating the need for single cell cloning to validate sgRNAs. Because Ba/F3 cells are a murine cell line, they are ideal for testing sgRNAs targeting the mouse genome that can subsequently be used for downstream *in vivo* applications. This system also allows for rapid and efficient testing of different permutations of sgRNA pairs with limited cloning steps, which is especially useful when generating fusions.

**Figure 1.**
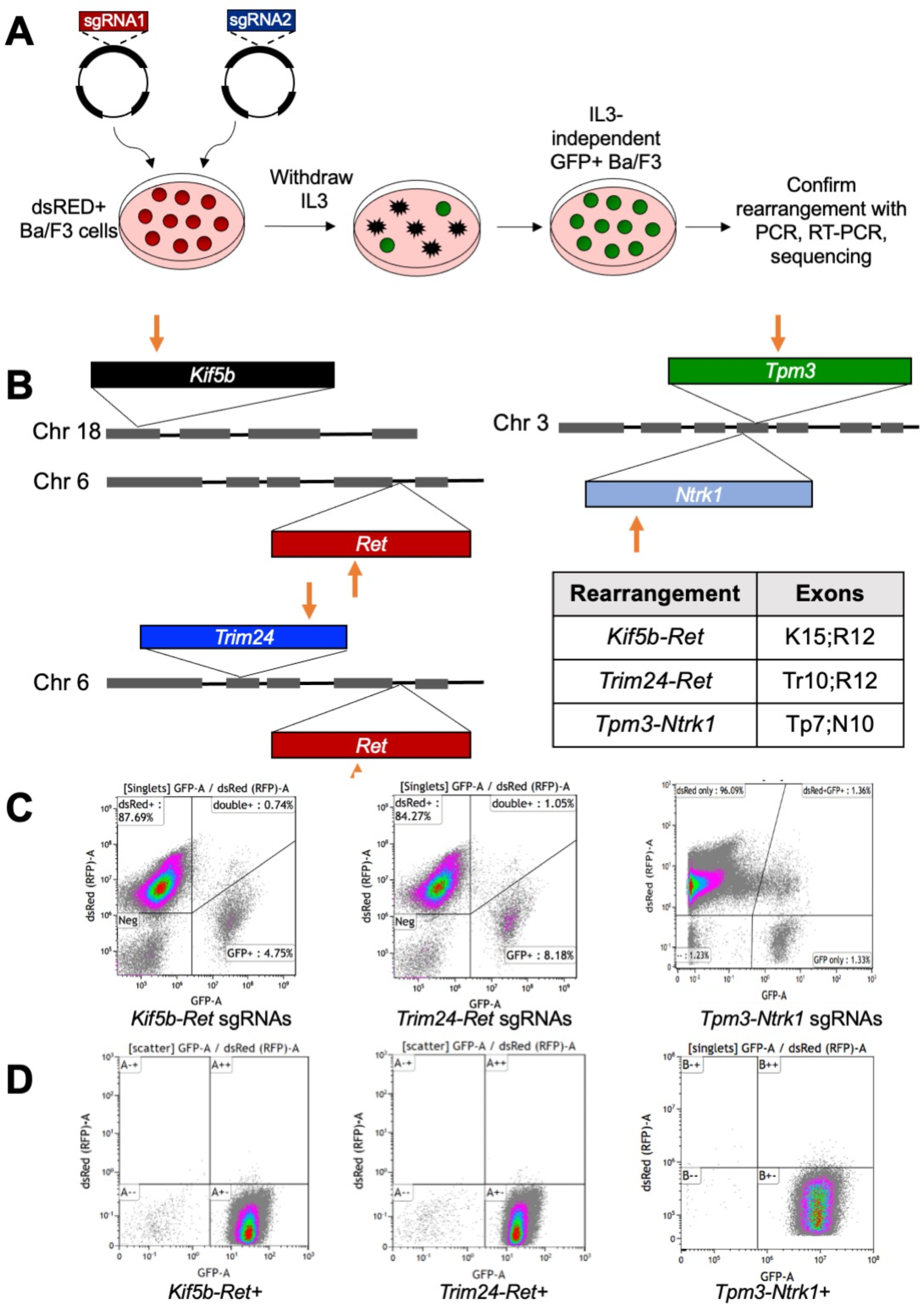
Ba/F3 based sgRNA screening technique. **A.** Schematic describing sgRNA screening technique in Ba/F3 cells. **B.** Schematics of CRISPR/Cas9 induced chromosomal rearrangements in *Ret* and *Ntrk1*. Red arrows indicate approximate genomic loci targeted by sgRNAs. Table of specific exons included in *Ret* and *Ntrk* rearrangements (K, *Kif5b*; R, *Ret*; Tr, *Trim24*; Tp, *Tpm3*; N, *Ntrk1*)**. C-D.** Flow cytometry for dsRED and GFP expression in Ba/F3 dsRED+ cells 72 hours after transfection with sgRNAs (**C**) and after acquisition of IL3-independence (**D**).

In order to monitor transfection efficiency, we first transduced Ba/F3 cells with a floxed dsRED gene followed by an eGFP gene [31]. This results in a stable cell line labeled with dsRED until a Cre recombinase protein is expressed, which will excise the dsRED gene and allow transcription of eGFP yielding dsRED-, eGFP+ cells. In order to use GFP positivity as a surrogate for successful transfection, we engineered the pX330 Cas9-containing plasmid to also contain the Cre recombinase gene. After transduction with the conditional eGFP expression virus, we selected dsRED+ cells using FACS (**Supplementary Figure 2**). As a positive control, we tested this method with validated sgRNAs described by Maddalo et al. that generate *Eml4-Alk* rearrangements both *in vitro* and *in vivo* [26]. We transfected dsRED+ Ba/F3 cells with the pX330-*Alk-Eml4* plasmid containing both *Eml4* and *Alk* sgRNAs. Flow cytometry was used to assess transfection efficiency and presence of GFP+ cells (**Supplementary Figure 3A**). After approximately three weeks, IL3-independent cells were proliferating normally. We confirmed that IL3-independent clones harbored the *Eml4-Alk* rearrangement with genomic PCR (**Supplementary Figure 3B**). Ba/F3+*Eml4-Alk* cells were ∼8 times more sensitive to the ALK inhibitor crizotinib, than the parental Ba/F3 dsRED+ cell line supplemented with IL3 (**Supplementary Figure 3C**). Additionally, the resulting *Eml4-Alk*+ population was almost entirely GFP+ (**Supplementary Figure 3D**). These findings confirm that the sgRNA screening method successfully selects for cells that have acquired the intended oncogenic rearrangement.

A central goal of this study was to generate novel *Ret* and *Ntrk1* cancer models. These oncogenes are observed in NSCLC and other cancer types, but limited *RET*+ and *NTRK*+ human cell lines exist [5, 25, 32-34]. We designed sgRNAs predicted to generate the murine equivalents of three distinct fusion oncogenes, *Kif5b-Ret*, *Trim24-Ret* and *Tpm3-Ntrk1* (**Figure 1B**). We established that the dsRED+ Ba/F3 cells were successfully transfected with several combinations of sgRNA pairs, as assessed by flow cytometry for GFP+ cells (**Figure 1C**). In each case, we were successful in generating IL3-independent Ba/F3 cells in approximately 4-5 weeks which were nearly 100% GFP+ (**Figure 1D**). We believe that these cells took longer to attain IL3-independence than the *Eml4-Alk* cells because we transfected the sgRNA pair encoded on separate plasmids, while the *Eml4-Alk* sgRNAs were both encoded within a single plasmid. We confirmed that IL3-independent Ba/F3 cells harbored their intended chromosomal rearrangements with fusion-specific genomic PCR or RT-PCR (**Figure 2A**). Targeted sequencing of genomic DNA or cDNA further validated that *Kif5b-Ret*, *Trim24-Ret* and *Tpm3-Ntrk1* fusions were present (**Figure 2B**). It is interesting to note that with both *Ret* fusions, one copy of wild-type, non-rearranged *Ret* was preserved in the Ba/F3 cells (**Supplementary Figure 4**).

**Figure 2.**
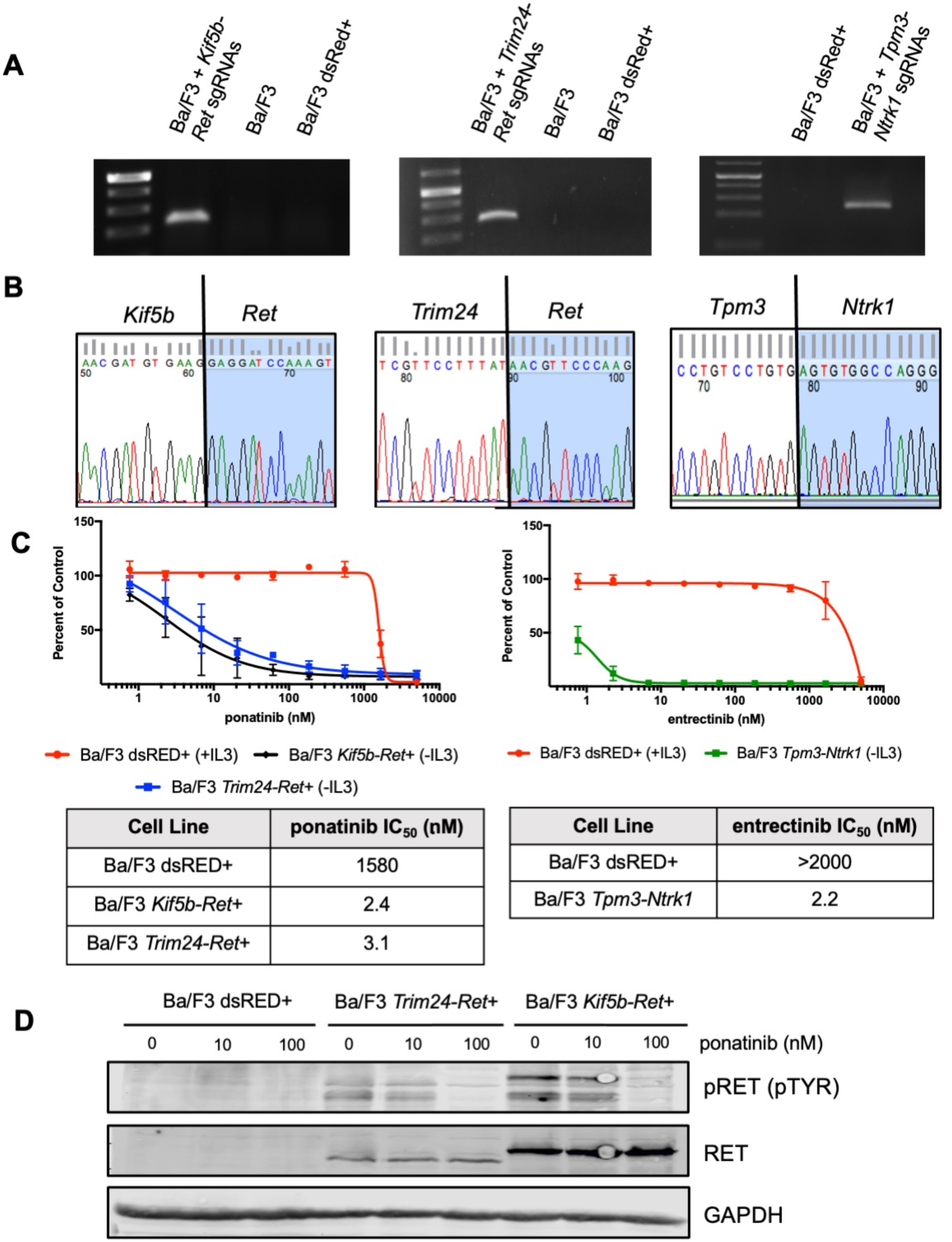
Ba/F3 cell-based screening identifies pairs of sgRNAs that successfully generate TKI-sensitive *Kif5b-Ret*, *Trim24-Ret* and *Tpm3-Ntrk1* rearrangements. **A.** Fusion specific RT-PCR for *Kif5b-Ret,* genomic PCR for *Trim24-Ret* or *Tpm3-Ntrk1*. **B.** Sequencing across fusion point in cDNA from Ba/F3 *Kif5b-Ret*+ cells and genomic DNA from *Trim24-Ret* and *Tpm3-Ntrk1.* C. (right) MTS proliferation assay performed on Ba/F3 dsRED+, Ba/F3 *Kif5b-Ret*+ and Ba/F3 *Trim24-Ret*+ cell treated with increasing doses of ponatinib. N=3 error bars represent ± SEM. (left) MTS proliferation assay performed on Ba/F3 dsRED+ and Ba/F3 *Tpm3-Ntrk1*+ cells treated with increasing doses of entrectinib. N=3 error bars represent ± SEM. The IC_50_ values are tabulated beneath the dose-response curves and were calculated with the Prism software program. **D.** Immunoblot analysis of Ba/F3 dsRED+, *Trim24-Ret*+ and Ba/F3 *Kif5b-Ret+* cells treated with ponatinib for 2 hours.

Finally, we were able to demonstrate that *Kif5b-Ret*+ and *Trim24-Ret*+ Ba/F3 cells were sensitive to multiple RET inhibitors, including ponatinib, cabozantinib, alectinib and foretinib (**Figure 2C, Supplementary Figure 5A-D**). We were also able to detect the expression of the RET fusion protein in *Trim24-Ret*+ and *Kif5b-Ret*+ Ba/F3 cells (**Figure 2D**). As expected, phosphorylation of RET could be inhibited upon treatment with ponatinib, a pan-TKI with significant RET activity (**Figure 2D**). Similarly, *Tpm3-Ntrk1*+ Ba/F3 cells were significantly more sensitive to the TRK inhibitor entrectinib than parental Ba/F3 sdRED+ cells (**Figure 2C**). Collectively, these data demonstrate that this method can efficiently screen and enrich for pairs of sgRNAs that generate distinct and functional fusion oncogenes.

To test the ability of the validated pairs of *Trim24* or *Kif5b*-*Ret* sgRNAs to induce lung tumors in mice, the sgRNA pairs shown in **Supplementary Figure 1** were cloned into the Ad-Cas9-2A-Cre adenoviral vector encoding Cas9 to catalyze the intended CRISPR-dependent gene rearrangements and Cre recombinase to induce LoxP-mediated excision of floxed alleles. Six weeks after instillation of Adeno-Cas9-Cre-*Trim24/Ret* gRNA virus in C57BL/6-TP53^fl/fl^ mice, multiple tumor foci were observed in the fixed lung lobes (**Figure 3A**), demonstrating that the *Trim24* and *Ret* sgRNAs induced gene rearrangement-dependent tumorigenesis as previously observed with *Eml4-Alk* rearrangements [11, 26]. Similarly, we introduced an adenovirus expressing the *Kif5b* and *Ret* sgRNAs into lungs of C57BL/6 mice without floxed TP53 and observed multiple tumors 10 weeks after instillation of adenovirus (**Figure 3B**). Thus, the *Trim24-Ret* and *Kif5b-Ret* rearrangements efficiently induce lung tumors in mice.

**Figure 3.**
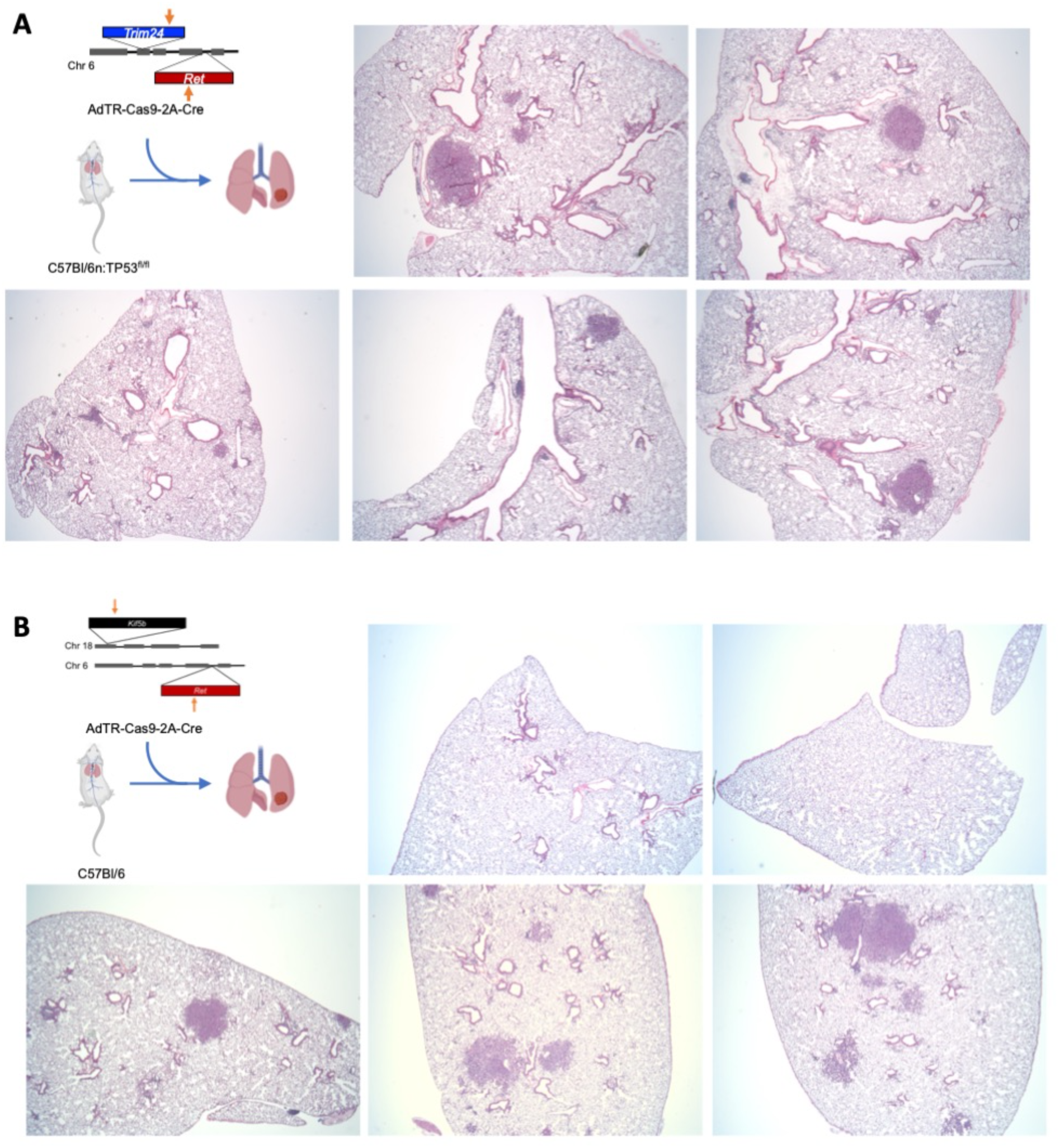
Murine model of TRIM24-RET fusion lung cancer. Design schema of CRISPR/Cas9 strategy to induce rearrangement at the (**A**) *Trim24* and *Ret* gene locus and (**B**) *Kif5b* and *Ret* for the generation of *Trim24-Ret* or *Kif5b-Ret* fusion respectively. **A.** TP53^fl/fl^ mice were intratracheally instilled with AdTR-Cas9-2A-Cre adenovirus and after 6 weeks, lung lobes were harvested, formalin-fixed, paraffin-embedded and sections were stained with H&E to assess histology. **B.** C57BL/6 mice were intratracheally instilled with Ad-TR-Cas9-2A-Cre adenovirus and after 10 weeks lung lobes were harvested, formalin-fix, paraffin-embedded and sections were stained with H&E.

To develop a murine *Trim24-Ret*-driven lung adenocarcinoma cell line, tumor-bearing lungs were harvested, minced and submitted to standard tissue culture techniques. A stable cell line was subsequently established, TR.1, that exhibited potent *in vitro* growth inhibition (**Figure 4A**) in response to the RET-specific TKIs [35], LOXO-292 (selpercatinib) and BLU-667 (pralsetinib). Immunoblot analyses of extracts from TR.1 cells treated for 2 hrs with *RET*-active TKIs (BLU-667, LOXO-292 and RXDX-105) and the 3^rd^ generation EGFR inhibitor, osimertinib, as a negative control demonstrated that the three *RET* inhibitors inhibited in a dose-dependent fashion levels of phospho-Y1062 RET, phospho-S473-AKT and phospho-ERK while osimertinib was without effect (**Figure 4B**). The TR.1 cell line was used to establish orthotopic tumors in the left lung lobe of syngeneic C57BL/6 mice (see Materials and Methods and [11, 12]). Following tumor establishment for ∼10 days, the initial tumor volumes were measured by μCT and the mice were treated with LOXO-292 (10 mg/kg) or diluent by oral gavage. Compared to diluent control, tumor shrinkage was induced by LOXO-292 treatment after 6 (31.4 + 9.1%) and 13 (27.4 + 5.2%) days of treatment (**Figure 5**). Within three weeks of LOXO-292 treatment, the TR.1 tumors began to progress, albeit at a slower rate than that exhibited by untreated tumors. The studies demonstrate that the TR.1 cell line is suitable for pursuing a variety of pre-clinical studies using an orthotopic implantation model.

**Figure 4.**
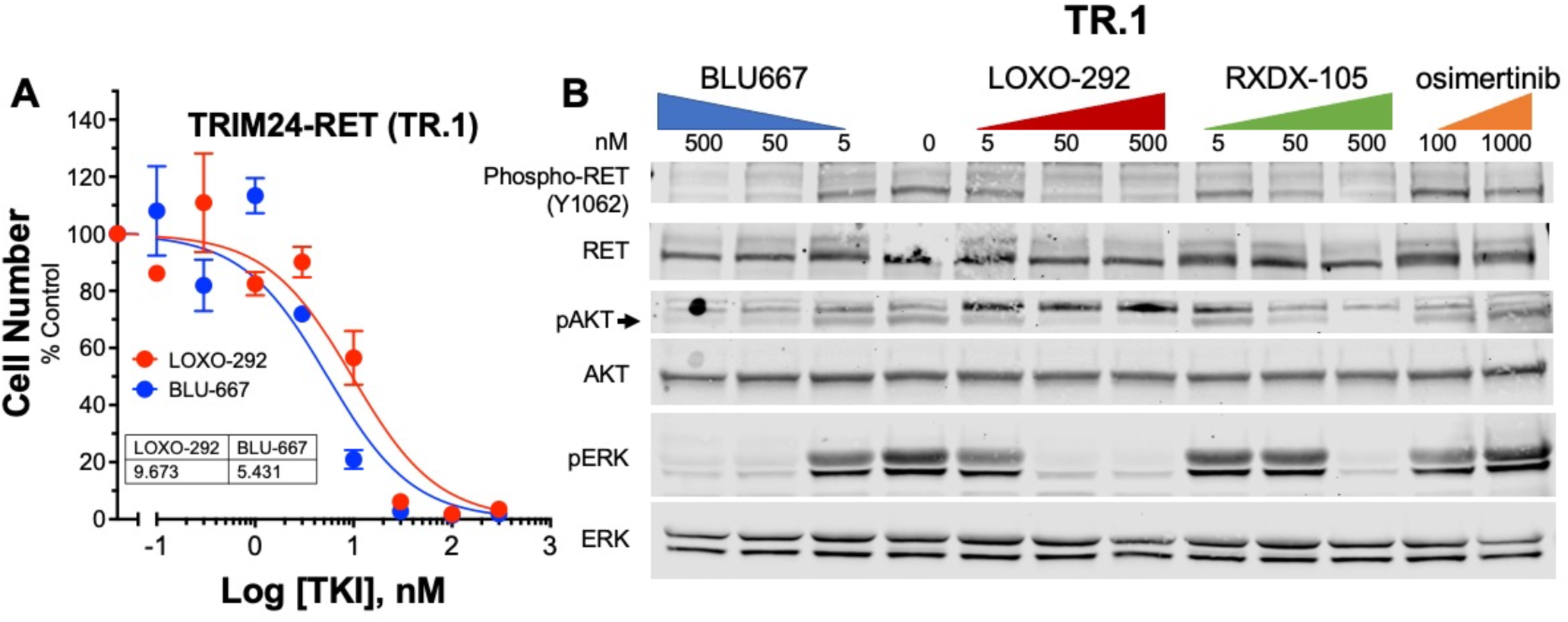
Derivation of a novel murine *Trim24-Ret* fusion cell line. **A**. TR.1 cells were seeded at 200 cells/well in 96-well plates and after 24 hours, treated for 7 days with LOXO-292 or BLU-667 at the indicated concentrations. Cell number was determined with CyQUANT reagent as described in the Materials and Methods. **B**. TR.1 cells were treated for 2 hours with the indicated TKIs and cell extracts were submitted to immunoblot analysis for phospho-Y1062-RET, phospho-Ser473 AKT and phospho-ERK1/2 as described in the Materials and Methods. The filters were subsequently stripped and reprobed for total RET, AKT and ERK.

**Figure 5.**
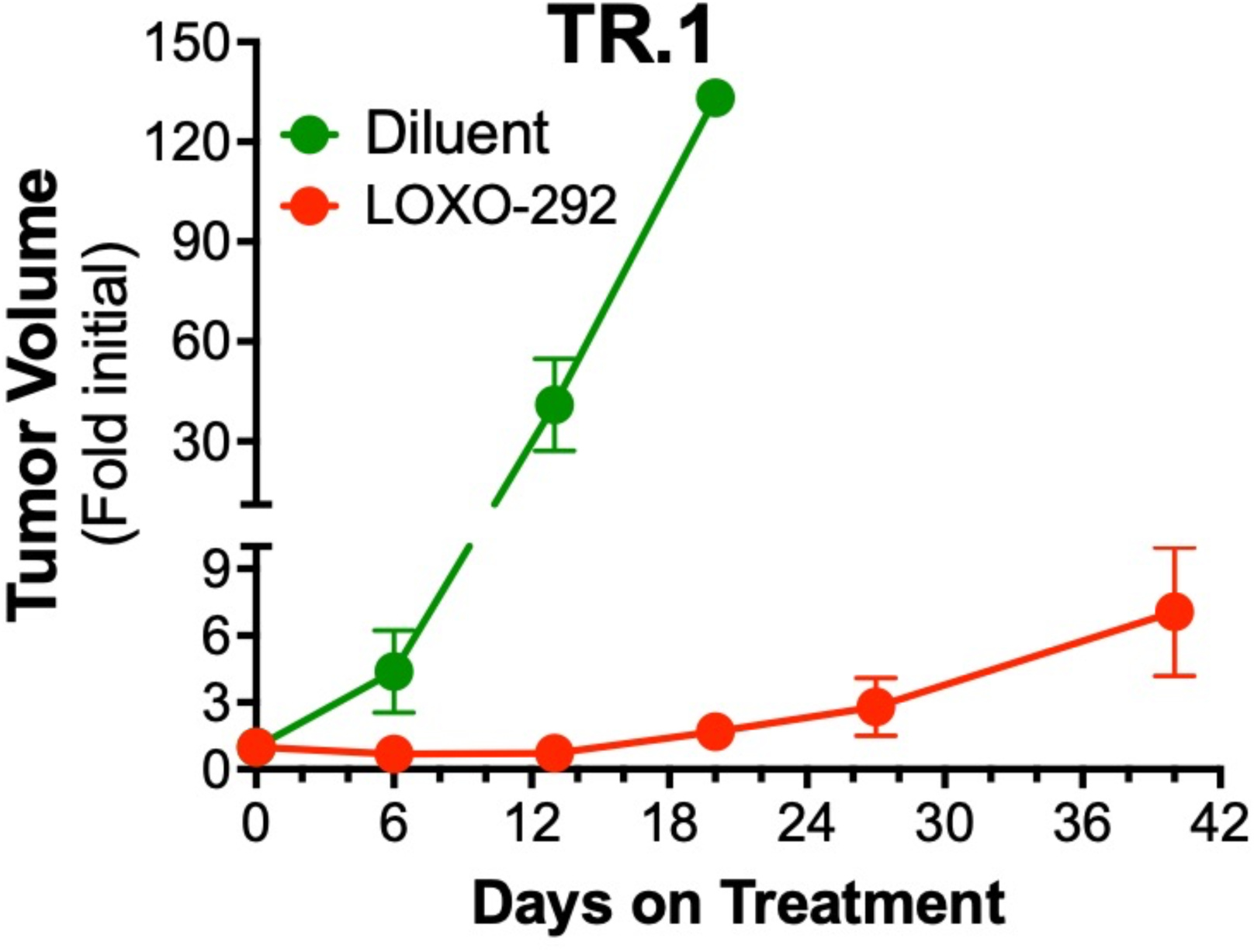
Sensitivity of TR.1 cell-derived orthotopic tumors to LOXO-292. TR.1 cells (500,000 cells per mouse) were implanted into the left lungs of C57BL/6 mice. The orthotopic tumors were allowed to establish for ∼10 days and the mice were submitted to μCT imaging to determine pre-treatment tumor volumes. Following randomization (n = 5 per group), the mice were treated daily with 10 mg/kg LOXO-292 or diluent by oral gavage. Mice were imaged by μCT weekly over the course of the experiments and tumor volume is presented as the fold change from the initial pre-treatment measurement. The initial tumor volumes (mean ± SEM) for the diluent and LOXO-292-treated groups were 2.4 ± 0.7 and 4.3 ± 91.2 mm^3^, respectively.

## Discussion

The Ba/F3 cell-based sgRNA screening method presented here allows for the validation of sgRNAs prior to dedicating time and resources to larger experiments or *in vivo* studies. We believe that this technique will be extremely versatile and expect that it can be used to select for other types of oncogenic alterations, such as point mutations or small InDels. In this regard, the design of sgRNA pairs that result in deletion of MET exon 14 to yield the “MET exon 14 skip” oncogene is predicted to be straightforward and yield murine models for this targetable oncogene. The generation of murine cell lines driven by *Kif5b-Ret* and *Tpm3-Ntrk1* fusion oncogenes is presently ongoing. This method poses some potential limitations, including the inability to screen interesting non-transforming alterations, and the possibility that Ba/F3 cell chromatin structure may limit accessibility to certain regions of the genome. We attempted, but failed to identify pairs of sgRNAs (including *Gopc-Ros1, Sdc4-Ros1* and *Ezr-Ros1*), to induce distinct *Ros1* fusion oncogenes with this method and are developing Cre-inducible *Cd74-Ros1* and *Ezr-Ros1*-transgenic mice as an alternative. We suspect that other genetic alterations and/or protein expression profiles not present in Ba/F3 cells may be required for some transforming events. Overall, this strategy provides a simple and efficient protocol for validating oncogenic sgRNAs that should accelerate functional characterization of many transforming alterations. Ultimately, the application of this CRISPR/Cas9 strategy will permit the development of mice and murine lung cancer cell lines driven by relevant oncogenes described in patients for which human cell line and PDX equivalents are not readily available. We have recently reported the development of murine *Eml4-Alk*-driven lung cancer cell lines that readily form orthotopic lung tumors in C57BL/6 mice [11]. Notably, the orthotopically-implanted tumors in syngeneic hosts exhibited profound and durable shrinkage upon alectinib therapy, but transient responses in immune-deficient mice, thereby demonstrating a role of adaptive immunity in the therapeutic response to TKIs. A similar requirement for adaptive immunity is observed with murine lung cancer cell lines driven by oncogenic *Egfr* transgenes [12]. In fact, there is a growing literature supporting the requirement for host immunity in the therapeutic responses of cancers to diverse oncogene-targeted agents [16, 17]. Thus, development and deployment of transplantable murine equivalents of oncogene-driven human cancers will permit a mechanistic investigation of these observations in fully immune-competent mice where similar studies using human cancer cell lines or PDXs will require immune-deficient mouse strains reconstituted with humanized immune systems.

Compared to the durable and profound TKI responses observed in mice bearing orthotopic *Eml4-Alk* and *Egfr^del19^* tumors [11, 12], the depth of response of the *Trim24-Ret* cell line, TR.1, to LOXO-292 described herein was quite modest (**Figure 5**). Moreover, clear evidence of tumor progression after ∼2 weeks of LOXO-292 therapy was observed, suggesting rapid acquisition of TKI resistance. In a recent study with murine *Egfr^del19^* cell lines, we explored mechanisms that may mediate rapid progression of the tumors under continuous osimertinib treatment when propagated in immune-deficient mice [12]. Induction of HGF and MET expression was observed in the progressing tumors and a cell line established from a TKI-resistant tumor exhibited acquired sensitivity to the MET inhibitor, crizotinib. As MET pathway activation is considered a *bona fide* bypass resistance pathway in human EGFR-driven lung cancers, murine oncogene-driven lung tumors propagated in immune-competent hosts may represent tractable model systems to explore acquired drug resistance and to define novel combination therapy strategies for prolonging progression free survival.

This approach also allows for efficient, simultaneous manipulation of other mutations, such as loss of tumor suppressors. There is a growing understanding that the genetic context in which driver oncogenes occur can substantially alter the tumor biology. The role of tumor suppressor loss in relationship to therapy response, development of resistance and interaction with the tumor microenvironment is in need of further investigation. In particular, it has been shown that tumor suppressors have differential ability to promote oncogenesis in the context of *KRAS* mutated lung cancer with distinct therapeutic vulnerabilities [36, 37]. We found that our *Trim24-Ret* rearrangement, which was introduced in mice bearing floxed *Tp53* genes, formed tumors much more rapidly than our *Kif5b-Ret* rearrangements in *Tp53* wild-type mice (6 weeks vs. 10 weeks). Indeed, this CRISPR-induced fusion method technique could be paired with novel technologies such as that employed by Cai et al. [37] to explore the functional impact of tumor suppressors on oncogenic RTKs without the need to generate genetically engineered mouse models for each oncogene.

## Author Contributions

LS, ATL and RCD conceived and designed the Ba/F3 screen studies, ATL developed the stable *Trim24-Ret* cell line (TR.1) and TKH, AN, RAN and LEH conceived and designed the orthotopic *in vivo* experiments. LS, ATL, TKH, AN and SKN performed the experiments presented herein. All authors contributed to interpretation of results as well as writing and editing the manuscript.

## Acknowledgements

We thank Ignyta for supplying entrectinib and Chugai Pharmaceuticals for providing alectinib. We would like to thank the University of Colorado Cancer Center Flow Cytometry Shared Resource and the Barbra Davis Center for Diabetes Molecular Biology Service Center for help with experiments.

