## Supplementary Information for "A Rapid, Functional sgRNA Screening Method for Generating Murine RET and NTRK1 Fusion Oncogenes"

| sgRNA Sequences | | |
|---|---|---|
| Rearrangement | sgRNA 1 (5' partner) | sgRNA 2 (3' partner) |
| *Kif5b-Ret* | 5'-GTTAAGTGAAAATCTTCAACG-3' | 5'-GCTATACCCACATAAGCCCCA-3' |
| *Trim24-Ret* | 5'-GATTGCTGAATAACCGCATAA-3' | 5'-GTTACCCTTGGGAACGTTACA-3' |
| *Tpm3-Ntrk1* | 5'-GTGCAAGTCTAGCATTAACAC-3' | 5'-GCTAGCTGGGACCCCGAAGTG-3' |

### Supplementary Figure 1

**Supplementary Figure 1.** Table of sgRNA sequences.

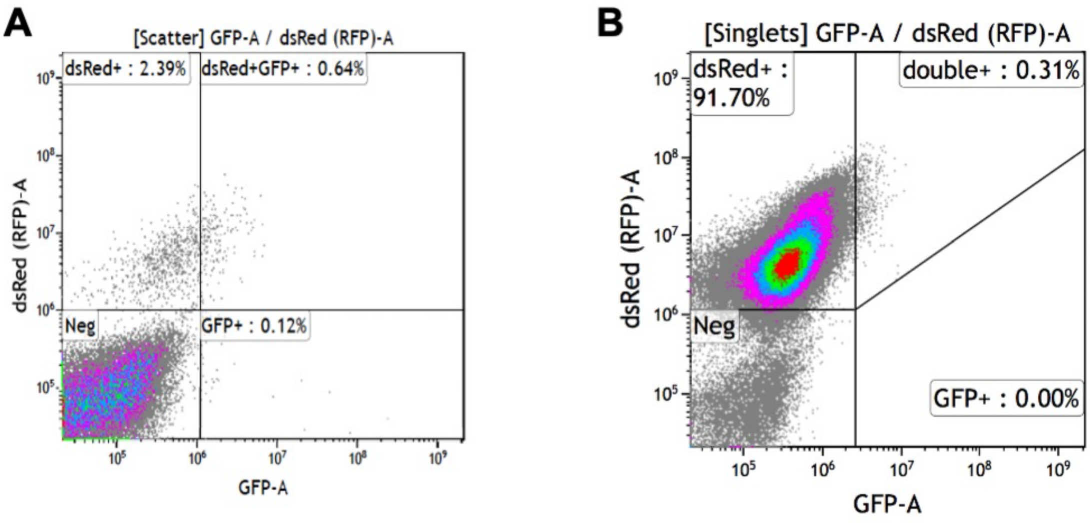


**Supplementary Figure 2**

**Supplementary Figure 2. Transduction and selection for dsRED+ cells. A.** Flow cytometry analysis for dsRED+ and eGFP+ Ba/F3 cells three days post transduction. **B.** dsRED+ Ba/F3 cell population after FACS.

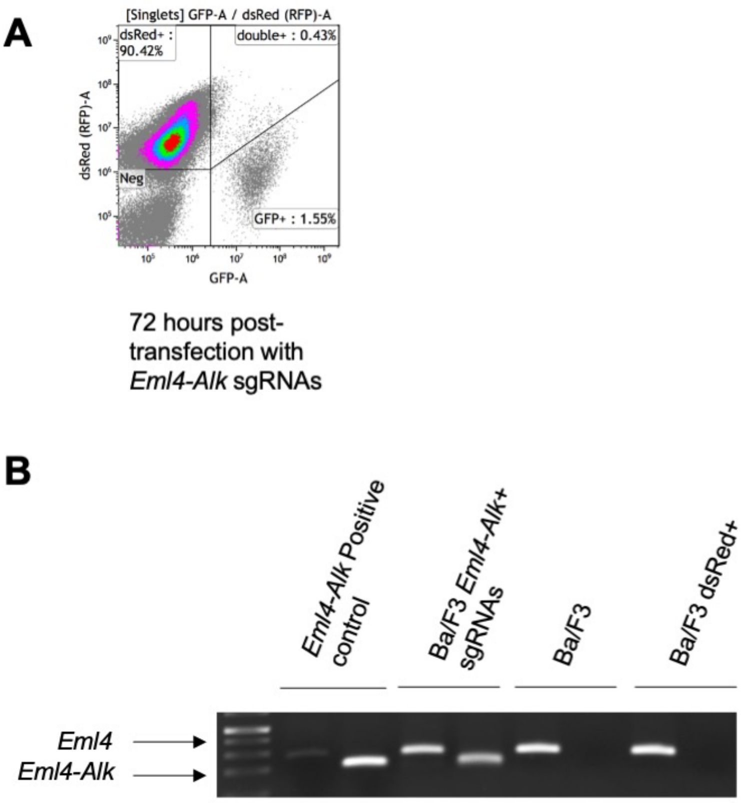


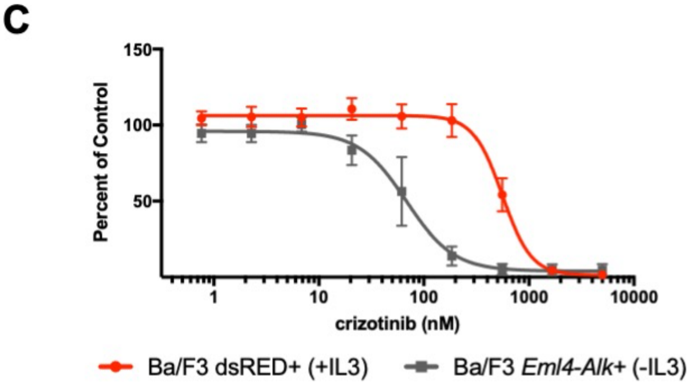


| Cell Line | crizotinib IC50 (nM) |
|---|---|
| Ba/F3 dsRED+ | 557.5 |
| Ba/F3 *Eml4-Alk*+ | 68.2 |

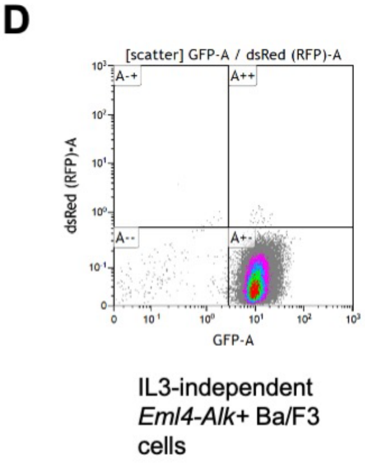


**Supplementary Figure 3**

**Supplementary Figure 3. Ba/F3 screening method selects and enriches for cells with CRISPR/Cas9 generated fusion oncogenes using validated *Eml4-Alk* sgRNAs. A.** Flow cytometry for dsRED and GFP after transfection with *Eml4-Alk* sgRNAs **B.** Genomic PCR using fusion specific primers, or non-rearranged *Eml4* primers in Ba/F3 + *Eml4-Alk* sgRNAs, parental Ba/F3 or Ba/F3 dsRED+ cells. "*Eml4-Alk* positive control" = DNA derived from mouse tumors confirmed to have *Eml4-Alk* rearrangements. **C.** MTS proliferation assay in Ba/F3 dsRED+ cells and Ba/F3 *Eml4-Alk*+ cells treated with increasing concentrations of crizotinib. **D.** Flow cytometry for dsRED and GFP after IL3 independence.

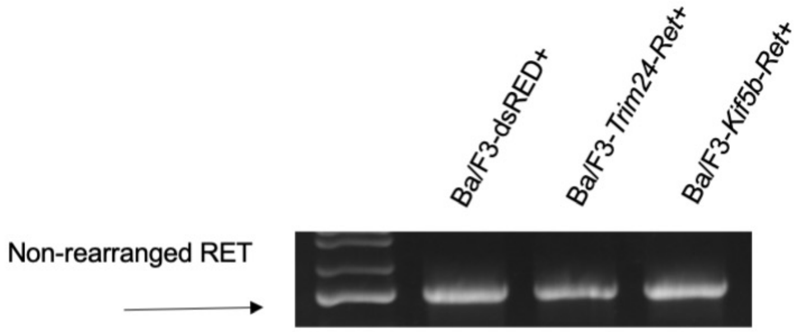


**Supplementary Figure 4**

**Supplementary Figure 4. Ba/F3 cells with *Kif5b-Ret* and *Trim24-Ret* rearrangements contain a copy of wild type, non-rearranged *Ret*.** Genomic PCR of *Ret* across intended fusion breakpoint.

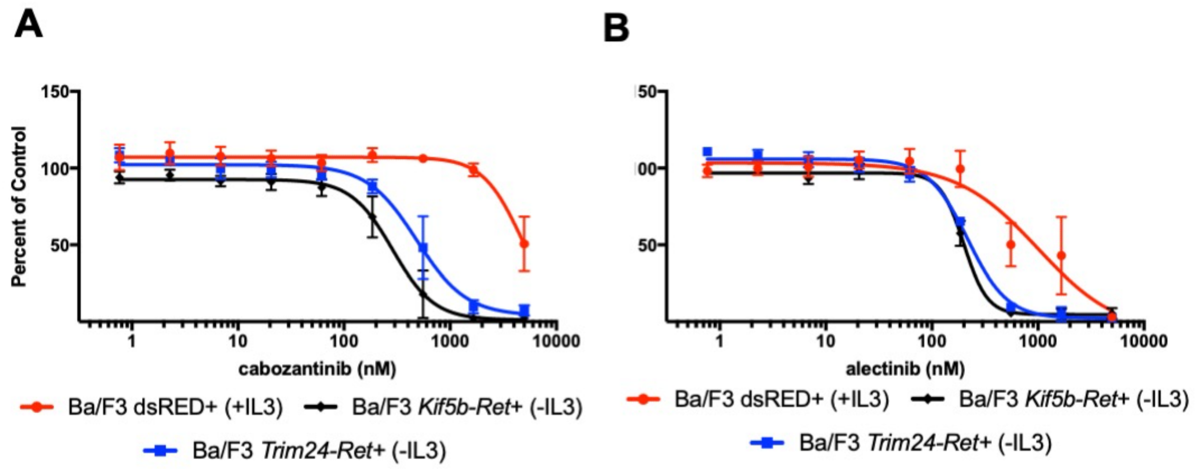


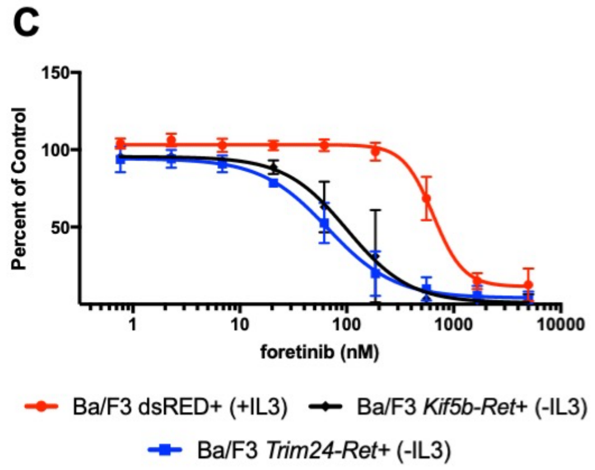


**D**

| Cell Line | Inhibitor | IC50 (nM) |
|---|---|---|
| Ba/F3 dsRED+ | cabozantinib | 4648 |
| Ba/F3 *Kif5b-Ret+* | | 287 |
| Ba/F3 *Trim24-Ret+* | | 484 |
| Ba/F3 dsRED+ | alectinib | 1000 |
| Ba/F3 *Kif5b-Ret+* | | 199 |
| Ba/F3 *Trim24-Ret+* | | 216 |
| Ba/F3 dsRED+ | foretinib | 647 |
| Ba/F3 *Kif5b-Ret+* | | 101 |
| Ba/F3 *Trim24-Ret+* | | 66 |

### Supplementary Figure 5

**Supplementary Figure 5. Ba/F3 cells with *Ret* rearrangements are sensitive to multiple RET inhibitors. A-C** MTS proliferation assay in cells treated with increasing concentrations of (**A**) cabozantinib (**B**) alectinib or (**C**) foretinib. N=3 error bars represent ± SEM. **D.** Chart of $IC_{50}$ values.
